# Found in Transcription: Gene fusions arise through defects in RNA processing in the absence of chromosomal rearrangements

**DOI:** 10.1101/825570

**Authors:** Yue Jiang, Michael J. Apostolides, Mia Husić, Robert Siddaway, Man Yu, Stephanie Mark, Arun K. Ramani, Cynthia Hawkins, Michael Brudno

## Abstract

Recent advancements in high throughput sequencing analysis have enabled the characterization of cancer-driving fusions, improving our understanding of cancer development. Most fusion calling methods, however, examine either RNA or DNA information alone and are limited to a rigid definition of what constitutes a fusion. For this study we developed a pipeline that incorporates several fusion calling methods and considers both RNA and DNA to capture a more complete representation of the tumour fusion landscape. Interestingly, most of the fusions we identified were specific to RNA, with no evidence of corresponding genomic restructuring. Further, while the average total number of fusions in tumour and normal brain tissue samples is comparable, their overall fusion profiles vary significantly. Tumours have an over-representation of fusions occurring between coding genes, whereas fusions involving intergenic or non-coding regions comprised the vast majority of those in normals. Tumours were also more abundant in unique, sample-specific fusions compared to normals, though several fusions exhibited strong recurrence in the tumour type examined (diffuse intrinsic pontine glioma; DIPG) and were absent from both normal tissues and other cancers. Intriguingly, tumours also show broad up- or down-regulation of spliceosomal gene expression, which significantly correlates with fusion number (p=0.007). Our results show that RNA-specific fusions are abundant in both tumour and normal tissue and are associated with spliceosomal gene dysregulation. RNA-specific fusions should be considered as a potential mechanism that may contribute to cancer formation initiation and maintenance alongside more traditional structural events.

## Introduction

Gene fusions represent an important class of genomic alterations. Fusion events can lead to increased oncogene expression, decreased expression of tumour suppressors, and formation of fusion proteins with oncogenic functions. The impacts of such changes are well documented in the tumourigenesis of multiple cancers (Mertens et al. 2015; Mitelman et al. 2007; Xiao et al. 2018; Yoshihara et al. 2015), with estimates suggesting that fusions account for nearly 20% of human cancer morbidity (Mitelman et al. 2007). They have thus been classically associated with the development of cancer, and the recurrence of specific fusions in some cases can act as a tumour-specific signature. Expression of fusion transcripts is typically taken to indicate chromosomal rearrangement, however RNA-specific fusions have been previously found in non-cancer cells and tissues (Babiceanu et al. 2016). Some recurrent fusions in normal tissues also show evidence of functional roles, challenging the notion that these are predominantly cancer-specific events (Li et al. 2008).

The detection of gene fusions has been greatly improved by recent advancements in high throughput sequencing (HTS). Approaches using multiple alignment steps for RNAseq analysis, such as EricScript, have led to the discovery of fusions associated with lung cancer (Benelli et al. 2012; Saber et al. 2016). deFuse allows for the identification of fusions at all possible locations within a genome, as opposed to solely at exon boundaries (McPherson et al. 2011). FusionMap introduced the use of junction-spanning reads to determine fusion junctions with base pair-level accuracy in either RNAseq or DNAseq data (Ge et al. 2011). Coupling of whole genome sequencing (WGS) and RNAseq provides especially strong evidence for fusion events, and has led to the discovery of oncogenic fusions in liver (Shiraishi et al. 2014), bladder (Guo et al. 2013) and lung cancers (Ju et al. 2012), as well as various other tumour types (Yoshihara et al. 2015). Recent tools, such as INTEGRATE, can examine WGS and RNAseq individually or in tandem (Zhang et al. 2016).

To study the nature of gene fusions in cancer, we designed a pipeline that incorporates the above four fusion callers. As individual callers can have biases towards specific categories of fusions, integrating multiple fusion-calling methods allows for detection of a broader variety of fusions. Our pipeline combines the results of these callers and can examine WGS data to determine if DNA support is present for fusions, which can yield insights into particularly complex cancers. With this, we examined RNAseq and WGS data of brain tissue samples from patients with diffuse intrinsic pontine glioma (DIPG), an aggressive childhood brainstem tumour with limited treatment options and a median survival of less than 1 year (Buczkowicz et al. 2014a). We also extended our analysis to subsets of normal tissue samples from the Genotype-Tissue Expression (GTEx) database (Lonsdale et al. 2013) and samples from various cancer types in The Cancer Genome Atlas Portal (TCGA) (The Cancer Genome Atlas Research Network 2008).

Our study provides novel insight into the nature of the fusions found in the transcriptomes of DIPG patients. Many of the fusions we identified, including ones that were successfully validated with RT-PCR, did not have genomic support from WGS sequencing. Most fusions specific to tumours were non-recurrent and occurred between coding genes. Moreover, DIPG tumour samples showed differential expression of spliceosomal genes, which correlated positively with fusion count and suggests that gene fusions in DIPG might be driven in part by transcriptional and splicing dysregulation. Our work expands the diversity of mechanisms driving fusion formation, suggesting that RNA-based fusions are an important component of the diversity in DIPG, and potentially in other tumours.

## Results

### Preliminary RNAseq and WGS analysis of DIPG samples

For this study, we obtained 34 DIPG tumour (DIPG-T) and 17 normal brain tissue (DIPG-N) samples from autopsies of 36 DIPG patients at the Hospital for Sick Children, with approval from the hospital Research Ethics Board. 15 DIPG-T samples and 15 DIPG-N samples comprised paired sets from the same 15 patients. We performed transcriptome sequencing for all samples. Genome sequencing was performed for 27 of these samples (14 DIPG-T and 13 DIPG-N) and published previously (Buczkowicz et al. 2014b). Initial RNAseq analysis of the DIPG-T samples using the fusion caller deFuse (McPherson et al. 2011) revealed a fusion between the epidermal growth factor receptor gene (*EGFR*) and the histone deacetylase 9 gene (*HDAC9*) gene in sample DIPG18T (Figure 1). deFuse reported over 400 split reads supporting this fusion with high mappability scores for the sequences surrounding the breakpoints. We subsequently validated its presence in RNA using RT-PCR and Sanger sequencing. Comparing *EGFR* and *HDAC9* gene orientations and the directions of their supporting read pairs, we found that the fusion is composed of the *EGFR* sense strand and *HDAC9* antisense strand.

**Figure 1.**
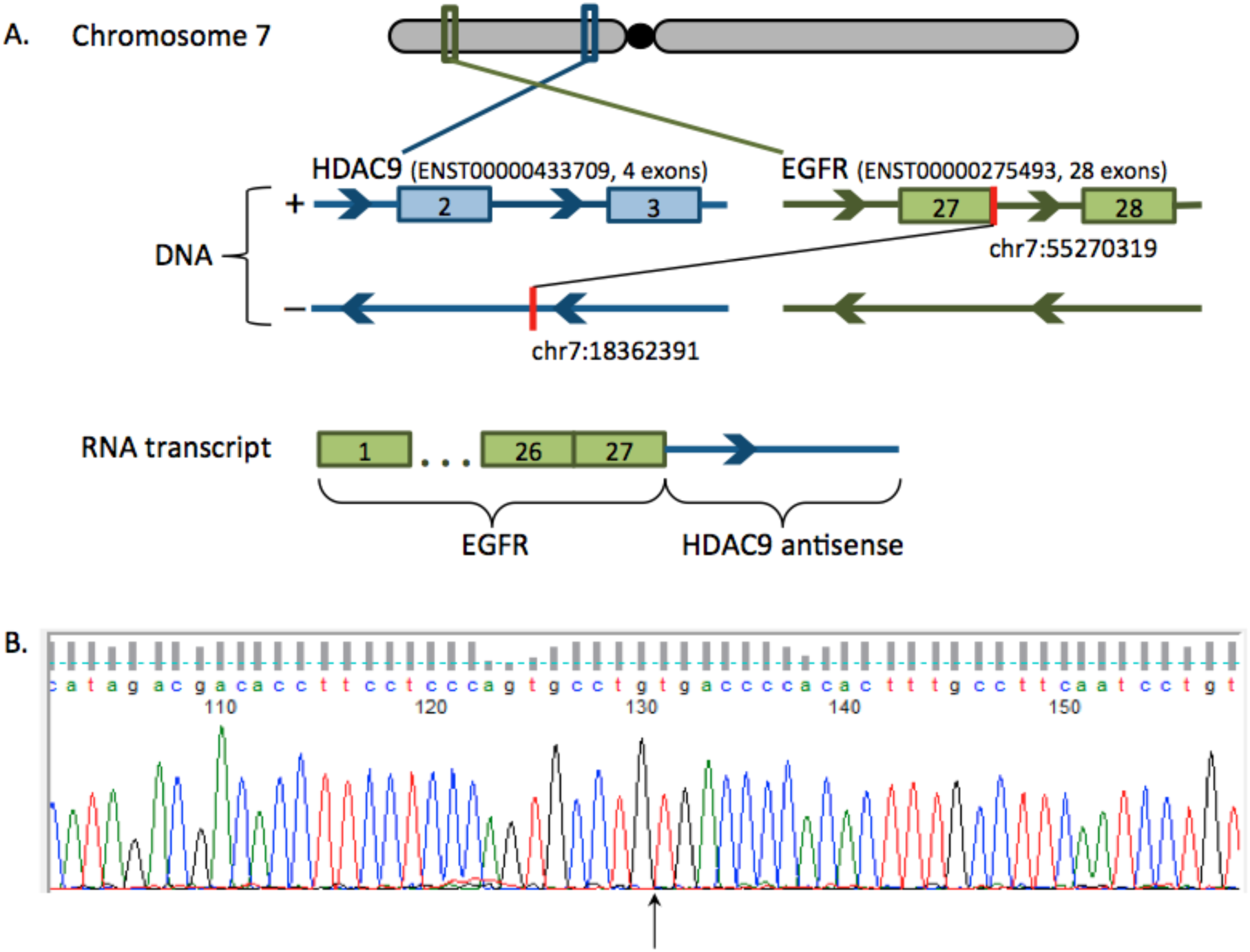
Illustration of the *EGFR-HDAC9* fusion in sample DIPG18T. (**A**) Structure of the fusion between the *EGFR* and *HDAC9* genes in DIPG18T. The fusion could only be identified using the fusion caller deFuse, and examination of the supporting reads revealed that it consists of the *EGFR* sense and *HDAC9* antisense strand. (**B**) Sanger sequencing of the DIPG18T transcriptome showed that *EGFR-HDAC9* is expressed. The sequencing trace is shown and the fusion junction indicated by the arrow.

To further confirm this fusion computationally, we analyzed the RNAseq data of DIPG18T using 3 additional fusion callers; however, none identified *EGFR-HDAC9*. Of all four fusion callers used, deFuse is the only one that reports fusions in intergenic regions, making it possible that the additional 3 callers simply dismissed *EGFR-HDAC9* due to the involvement of the *HDAC9* antisense strand.

Intriguingly, we found no DNA evidence for *EGFR-HDAC9* in the WGS data of DIPG18T. This was despite rigorous bioinformatic and manual analysis and multiple (genomic) PCR attempts to amplify the region using various experimental conditions with RNA-based primers, on the assumption that the introns are not long enough to evade PCR. This indicated that the fusion is not likely to be caused by underlying DNA structural variation, and led us to further investigate the presence of fusions in the transcriptomes of DIPG patients and if they have support in the DNA.

### Characterization of fusions in DIPG tumours compared to normal brain tissues from DIPG patients

Our *EGFR-HDAC9* results demonstrate that not all callers are able to equally identify fusions with specific characteristics. Therefore, to obtain a more complete picture of the fusion landscape in DIPG, we developed a fusion detection pipeline that consolidates results from multiple fusion callers (see Methods for details). As each caller has a unique methodology and definition of what constitutes a fusion, our pipeline allows for broader fusion detection than would be possible using just one method. This pipeline also uses a consistent annotation approach to name identified fusions based on specific categories as well as the distance and order of the constituent genes (Figure 2). It classifies fusions into 7 categories and filters them for uniqueness in mapping (BWAfilter), coverage (Covfilter) and supporting reads (Valfilter; all parameters described further in Methods).

**Figure 2.**
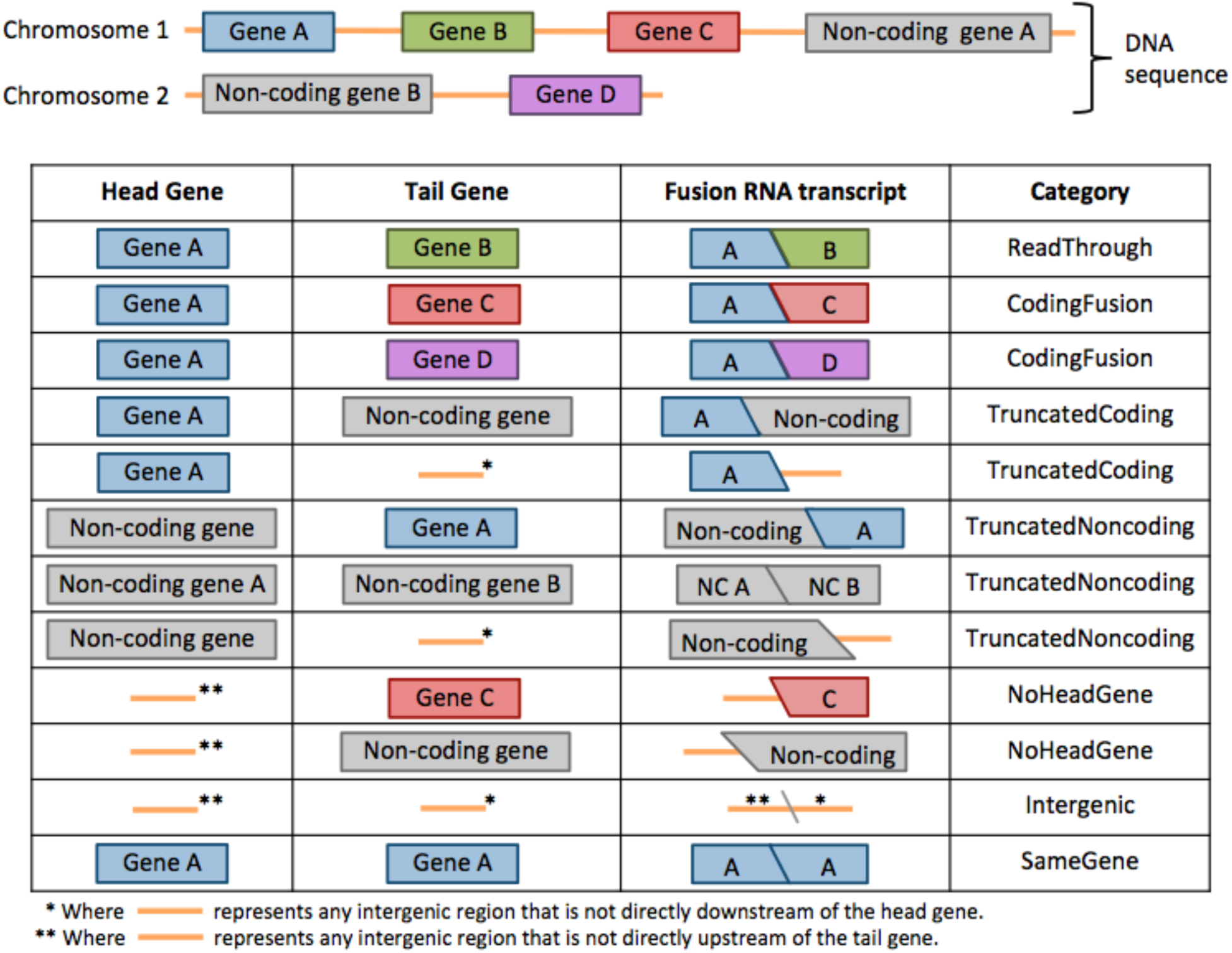
Depiction of the 7 categories used to distinguish fusions. Two chromosomes containing coding (coloured) and non-coding (grey) genes are shown above, with intergenic sequences represented by orange lines between genes. The chart below shows head and tail genes, the resultant fusion RNA transcript and the fusion category that the transcript would be assigned to. NC = non-coding.

#### Computational validation of the fusion detection pipeline

Prior to analyses of DIPG-T and DIPG-N samples, we evaluated the sensitivity and precision of our fusion detection pipeline using a previously published dataset of fusions in childhood glioma samples (Wu et al., 2014) as a validation set (Table 1, Supplementary Table S1). When a fusion from this validation set was required to have good mappability (BWAfilter30), at least 3 captured reads (Valfilter3) and at least 5 supporting reads (Covfilter5), our pipeline detected 90.9% of CodingFusions (50/55), 88.4% of TruncatedCoding (38/43) and 100% of TruncatedNonCoding (3/3) fusions. Except for one TruncatedCoding fusion detected by both deFuse and INTEGRATE, all TruncatedCoding and TruncatedNonCoding fusions were detected by deFuse alone. In previous works on fusion callers (Wu et al. 2013; Zhang et al. 2016; Okonechnikov et al. 2016) or the comparison of them (Liu et al. 2016), deFuse did not belong to the best performance group. This could be because most fusion callers do not report truncated fusions, leading to their exclusion from the validation benchmark. This would make it appear like deFuse’s false positive rate is much higher than it actually is.

**Table 1:** Fusion detection pipeline performance across different fusion categories.

|  | Wu et al. Validated +<br>BWAfilter30 | BWAfilter30 + Covfilter5 |  | BWAfilter30 +<br>Valfilter3 +<br>Covfilter5 |  |
| --- | --- | --- | --- | --- | --- |
| Category | Number of fusions | Sensitivity | Precision | Sensitivity | Precision |
| **CodingFusion** | 55 | 98.2% | 7.4% | 90.9% | 26.0% |
| **TruncatedCoding** | 43 | 93.0% | 8.5% | 88.4% | 12.4% |
| **TruncatedNoncoding** | 3 | 100.0% | 1.5% | 100% | 2.6% |
| **NoHeadGene** | 2 | 50.0% | 0.5% | 50.0% | 0.9% |
| **SameGene** | 1 | 0% | 0% | 0% | 0% |
| **All** | 104 | 94.2% | 4.3% | 88.5% | 11.1% |

Although the precision of our pipeline with respect to our validation dataset is low, it should be noted that every fusion in our pipeline’s output has at least 3 reads captured (Valfilter3). This suggests that all fusions reported by our pipeline are present in the RNAseq data, but are simply not included in the validation set. Overall, our results illustrate that our pipeline has high sensitivity in identifying fusions from RNAseq samples.

#### Occurrence of fusions in DIPG tumour samples and normal brain tissues

We first used the detection pipeline to analyze RNAseq data from the 15 paired DIPG-T and -N samples (n=30) for fusions that are specific to DIPG-T, specific to DIPG-N, or common to both sample types. We categorized the fusions and filtered them under the same parameters used for validation of the pipeline (BWAfilter30, Covfilter5, Valfilter3; Figure 3A, B).

**Figure 3:**
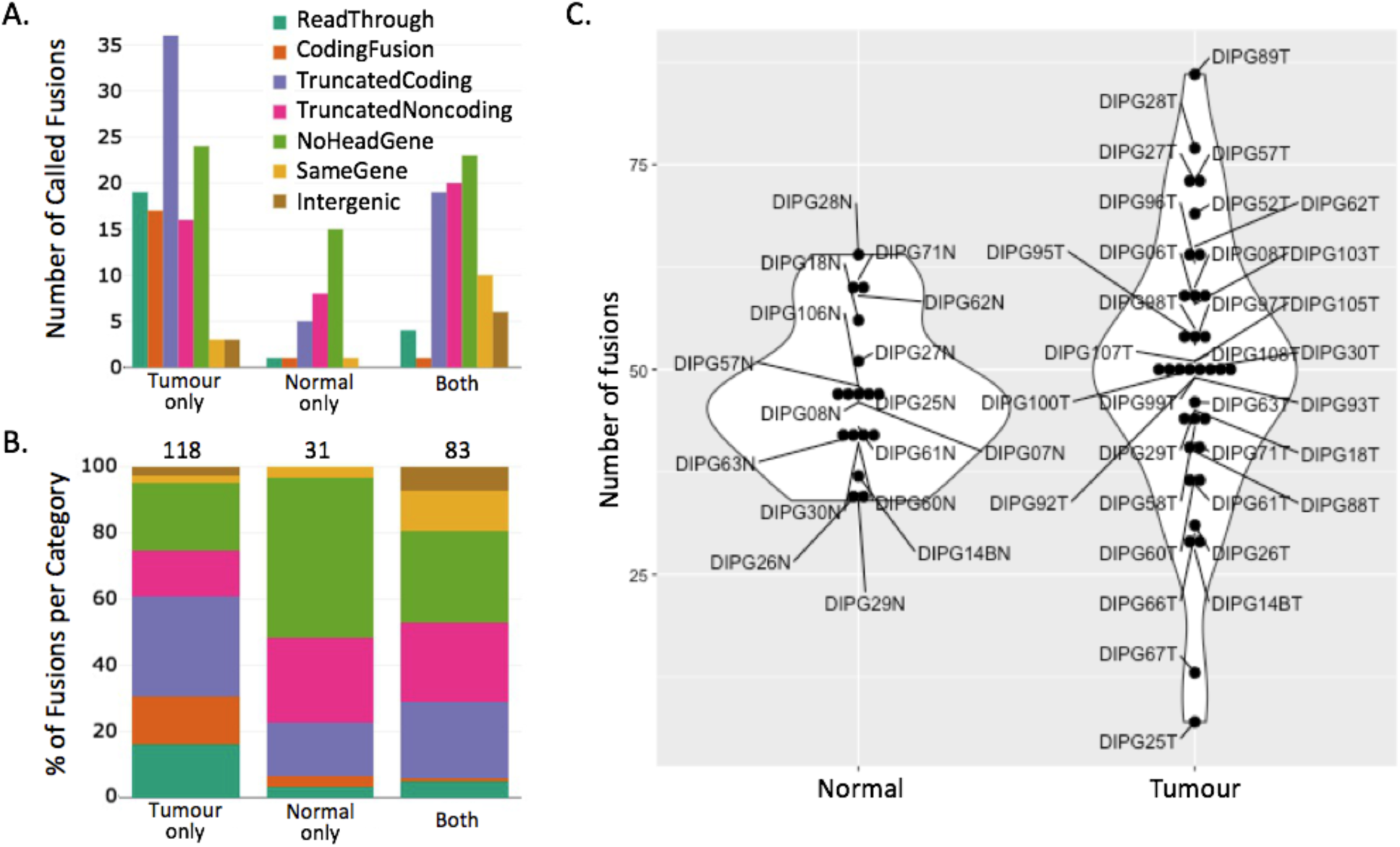
Distributions of fusions using BWAfilter30, Valfilter3 and Covfilter3 in paired (A,B) and unpaired (C) DIPG patient samples. (**A**) Number of fusions per fusion category in 15 DIPG-T/DIPG-N sample pairs. Counts are divided to represent fusions specific to DIPG-T samples (Tumour only), specific to DIPG-N samples (Normal only), or those common to both (Both). (**B**) The proportion of fusions from each category for each sample type in the 15 DIPG-T/DIPG-N samples pairs. Fusions between 2 coding genes appear to have a bias towards DIPG-T samples. For (A) and (B) the number of samples a given fusion is found in is not taken into account, only the sample type(s) it occurs in. (**C**) Distribution of total fusion counts for each of the 51 total DIPG-T and DIPG-N samples. While the average number of fusions per sample is similar between DIPG-T and DIPG-N, DIPG-T samples have a much larger variance in fusion number.

After filtering, we found a total of 201 fusions in DIPG-T samples and 114 fusions in DIPG-N samples. 232 unique fusions were present across all 30 paired samples --of these, 118 were DIPG-T-specific, 31 were DIPG-N-specific and 83 were present in both sample types (Figure 3A). Examining the fusion category proportions per sample type revealed that TruncatedCoding, TruncatedNoncoding and NoHeadGene fusions made up the majority of fusions in all sample groups (Figure 3B). They were particularly abundant in DIPG-N, making up 28/31 (90.3%) of DIPG-N-specific fusions in the paired sample set. SameGene fusions were most frequently found in both DIPG-N and DIPG-T. While their absolute numbers were relatively low, ReadThroughs and CodingFusions were much more likely to be DIPG-T-specific. CodingFusions in particular were almost entirely absent from DIPG-N, with 16/18 (88.8%) being unique to DIPG-T. Likewise, 19/24 (79%) ReadThroughs were DIPG-T-specific. These categories are the only ones representing fusions between 2 coding genes. We observed similar trends when extending this analysis to all 51 unpaired samples (Supplementary Figure S1).

As a given fusion may occur in multiple samples, we used 2 measures to calculate the average number of fusions per sample: 1) the average total number of fusions, which considers all fusions present in a given sample regardless of their occurrence in other samples, and 2) the average number of unique fusions, which considers the number of unique fusions that individual samples contribute to the pool of 232 total fusions. DIPG-T and DIPG-N had the same average total fusions per sample (33.4 and 33.7, respectively) yet differed in the average number of unique fusions per sample (13.4 and 7.6, respectively). Of note, the total number of fusions between individual DIPG-T samples varied widely compared to DIPG-N, which had a much tighter distribution of fusions per sample; again, this was true for both paired (Supplementary Figure S2) and unpaired (Figure 3C) sample sets.

#### Recurrence of fusions in DIPG-T and DIPG-N samples

Next, we examined the recurrence of fusions in DIPG and found that the vast majority of fusions that are recurrent in DIPG-T could also be found in DIPG-N (Supplementary Table S2). This was particularly true when requiring fusions to be found in 3 or more DIPG-T samples. This pattern was present across all fusion categories for the paired sample set (Supplementary Table S2a) and in all categories except ReadThroughs for the unpaired sample set (Supplementary Table S2b). These results indicate that fusions specific to DIPG-T are more likely to be unique to a single sample, while recurrent fusions are likely to be present in normal tissues as well as in tumours.

To confirm that fusions occur in normal tissues and that recurrent fusions present in both DIPG-T and DIPG-N samples are not due to contamination, we investigated the occurrence of CodingFusions in RNAseq data of normal brain tissue samples catalogued in the GTEx database. Our analysis revealed 51 CodingFusions in normal tissue GTEx samples, 4 of which were also found in DIPG-N samples (Supplementary Table S3). Finding such an abundance of fusions between coding genes in GTEx RNAseq data that pass strict filtering by our pipeline further indicates that fusions in normal tissues cannot be simply dismissed as false positives or contaminations.

### Identifying fusions specific to DIPG tumours

Our above results show that although many fusions in DIPG-T are sample-specific, several recurrent fusions are present as well. Such fusions may prove useful as biomarkers and provide insight into DIPG pathogenesis. Previous studies conducted at St Jude’s Children’s Hospital indicate that 5 fusions involving the 3 *NTRK* genes --which promote differentiation and survival of neuronal cells throughout the nervous system (Deinhardt and Chao 2014) --are associated with pediatric high-grade gliomas (pHGG) including DIPG (Wu et al. 2014). Two fusions (*BTBD1-NTRK3* and *VCL-NTRK2*) were present in DIPG, while the other 3 (*NTRK2-BEND5, ETV6-NTRK3, TPM3-NTRK1*) were specific to supratentorial pHGG (Wu et al. 2014). All five fusions were present in RNAseq data, however only three were also identified in corresponding WGS samples. Examining the St Jude’s RNAseq datasets with our own pipeline, we found that the five *NTRK* fusions passed all filters (Supplementary Table S4). We also detected a *BEND5-NTRK2* fusion that was not previously reported in the St Jude’s RNAseq data. This fusion has substantially different breakpoints from the reported *NTRK2-BEND5. NTRK* fusions have been previously associated with various tumours (Cocco et al. 2018), however both our pipeline and the analyses performed by Wu et al (2014) showed that the DIPG-specific *BTBD1-NTRK3* and *VCL-NTRK2* are unique to individual samples, and may serve as patient-specific signatures.

To identify which fusions in our sample cohort are exclusive to DIPG, we examined their occurrence in other tissue and cancer types. For this we developed a fusion validation pipeline that determines if a given fusion is present in an RNAseq dataset by examining that dataset for corresponding supporting reads (see Methods for details). Briefly, the validation pipeline aligns reads to both the putative fusion and to a combined annotation filter reference, and removes reads with stronger alignment to the filter reference, as these are likely false positives. We used this pipeline on two sample cohorts from the TCGA and GTEx databases to identify potential tumour-specific fusions and further refine those into fusions which are putative DIPG signatures. First we ran our pipeline (BWAfilter30, Valfilter3, Covfilter5) on our 51 unpaired DIPG-T and DIPG-N samples, and identified 430 fusions across all samples. From this, we selected a subset of 65 fusions present in 3 or more DIPG-T samples, and absent in DIPG-N samples (Supplementary Table S5). We used this as input to our validation pipeline (see Methods, Figure 8A-C) to query for these fusions in various tumour and normal samples from the TCGA and GTEx, as well as the DIPG/HGG dataset published by Wu et al. (2014). Finally, we calculated the fraction of samples in each tested sample group carrying each of the selected fusions (Figure 4).

**Figure 4.**
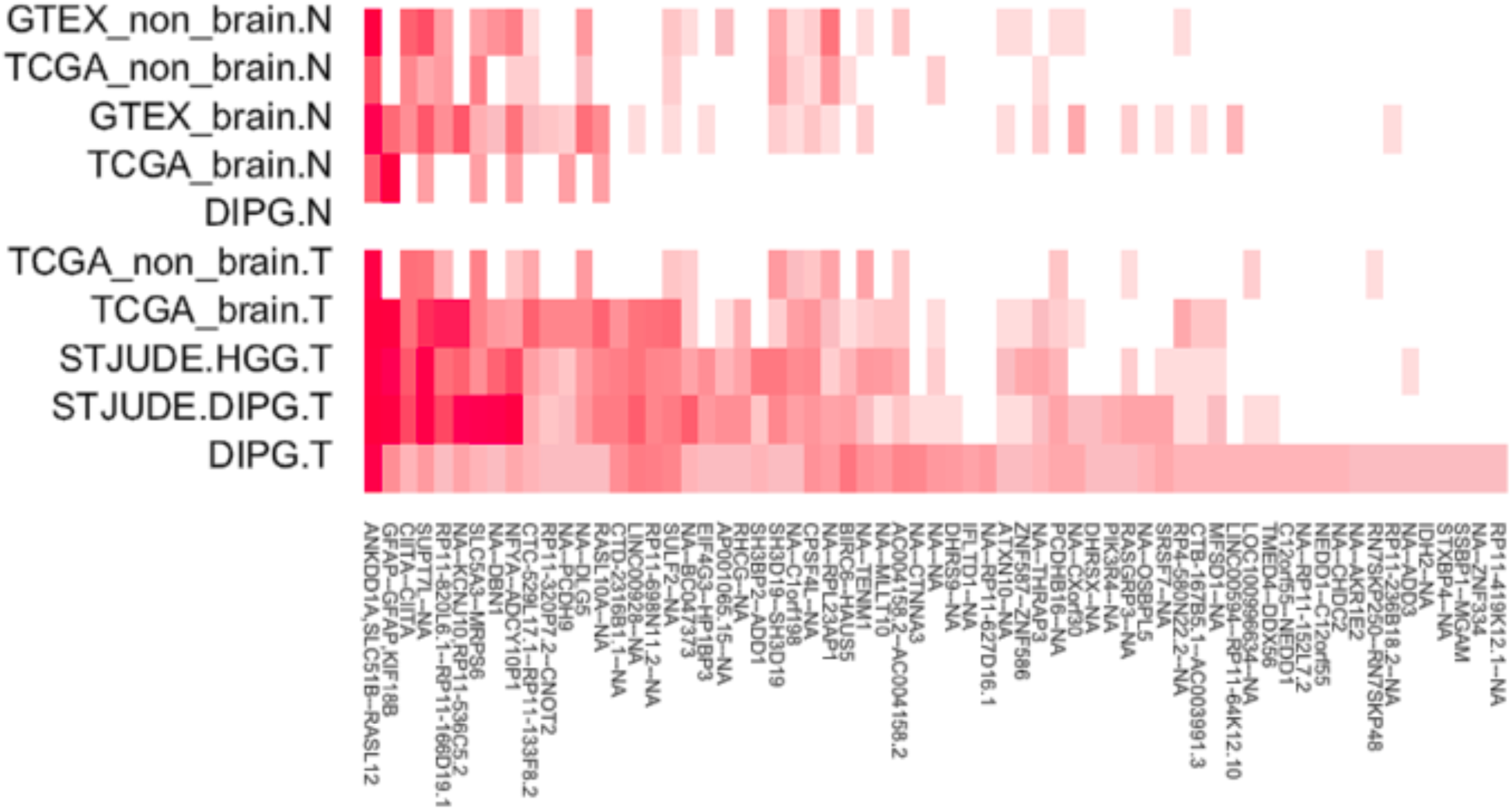
Heatmap of fusion expression across tumours and normal tissues. The expression of 65 DIPG-T-specific fusions (columns) was assessed in various TCGA brain and non-brain tumour samples, GTEx brain and non-brain normal tissue samples and DIPG and HGG samples from Wu et al 2014 (rows). Darkness of red corresponds to fraction of samples in each sample set containing a given fusion. T = tumour; N = normal; STJUDE = sample sets from Wu et al. 2014.

**Figure 5:**
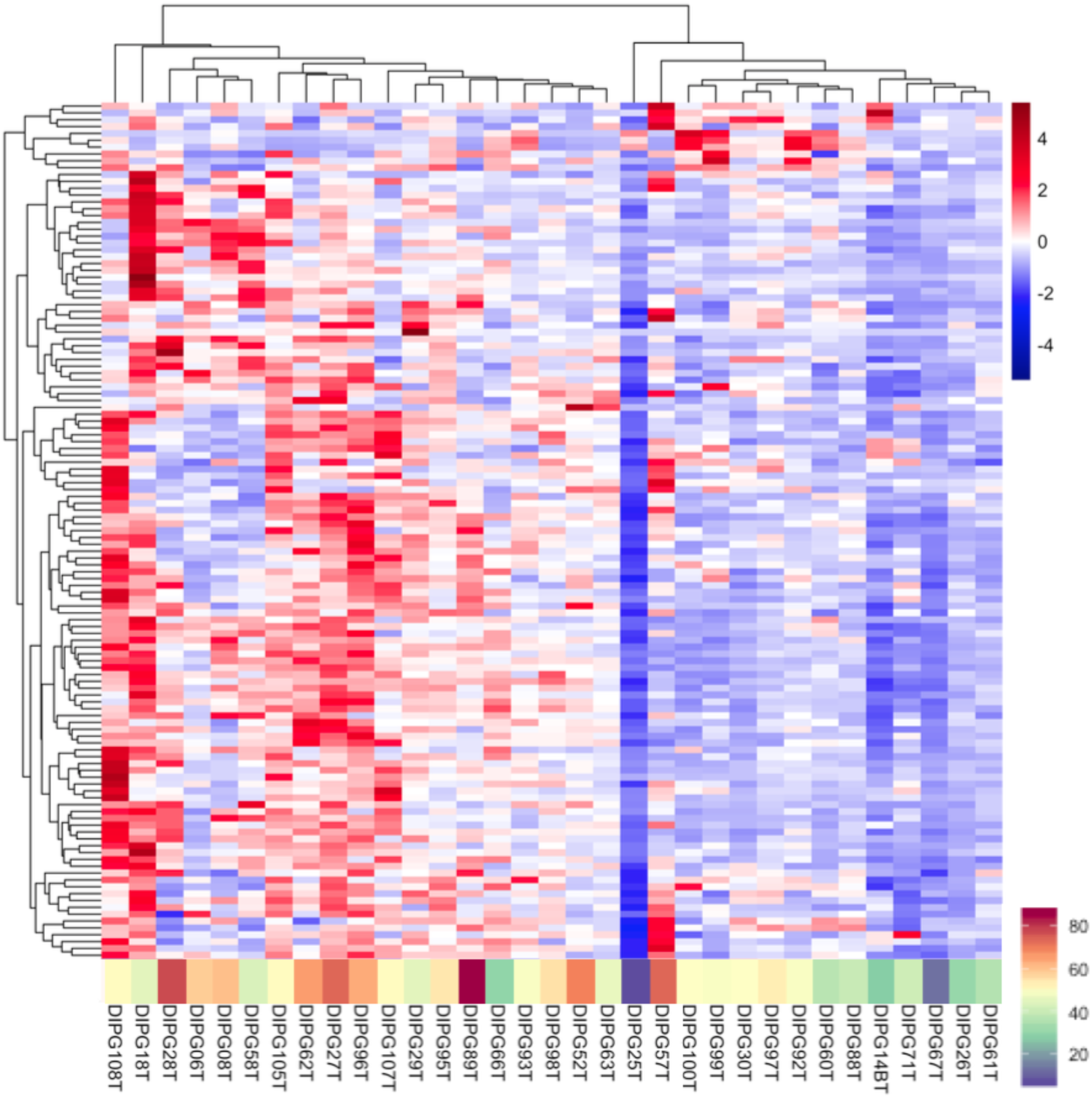
DIPG-T samples tend to broadly over- or under-express genes in the KEGG spliceosomal category. Heatmap: DIPG-T samples cluster into two groups showing either increased expression (left) or decreased expression (right) of 125 spliceosomal genes. Range of gene expression is shown in the scale on the top left where red indicates higher gene expression and blue indicates lower gene expression. Bottom panel: Measurement of the number of fusions in all categories per sample. DIPG-T samples that over-express spliceosomal genes typically contain a greater number of fusions, while samples under-expressing spliceosomal genes typically contain fewer fusions. The fusion count scale is depicted to the right, where the mean number of fusions in DIPG-N was used as the midpoint. Fusion counts were determined using the fusion detection pipeline with BWAfilter30, Valfilter3 and Covfilter5.

**Figure 6:**
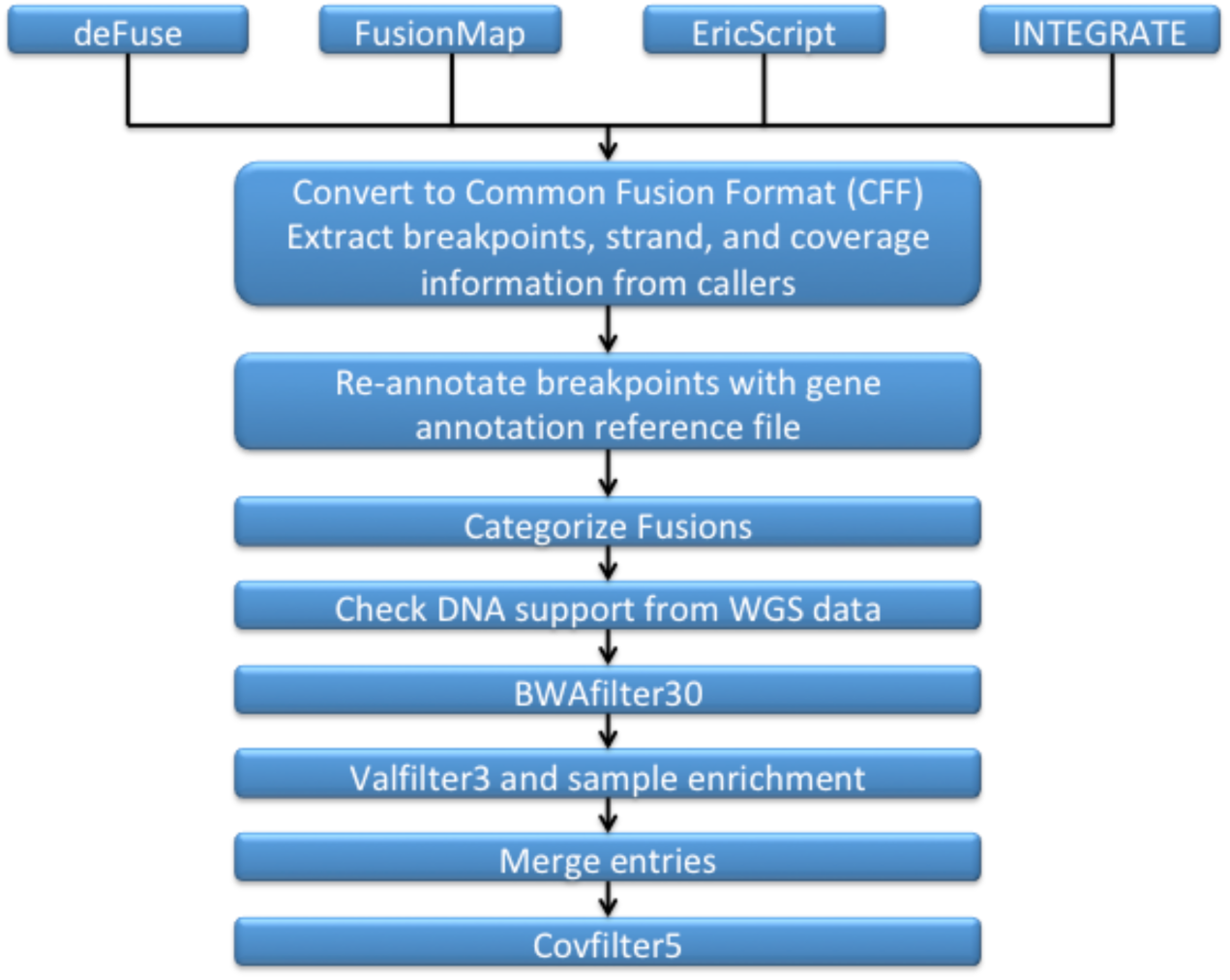
Workflow of the fusion detection pipeline used to identify fusions from RNAseq data.

**Figure 7.**
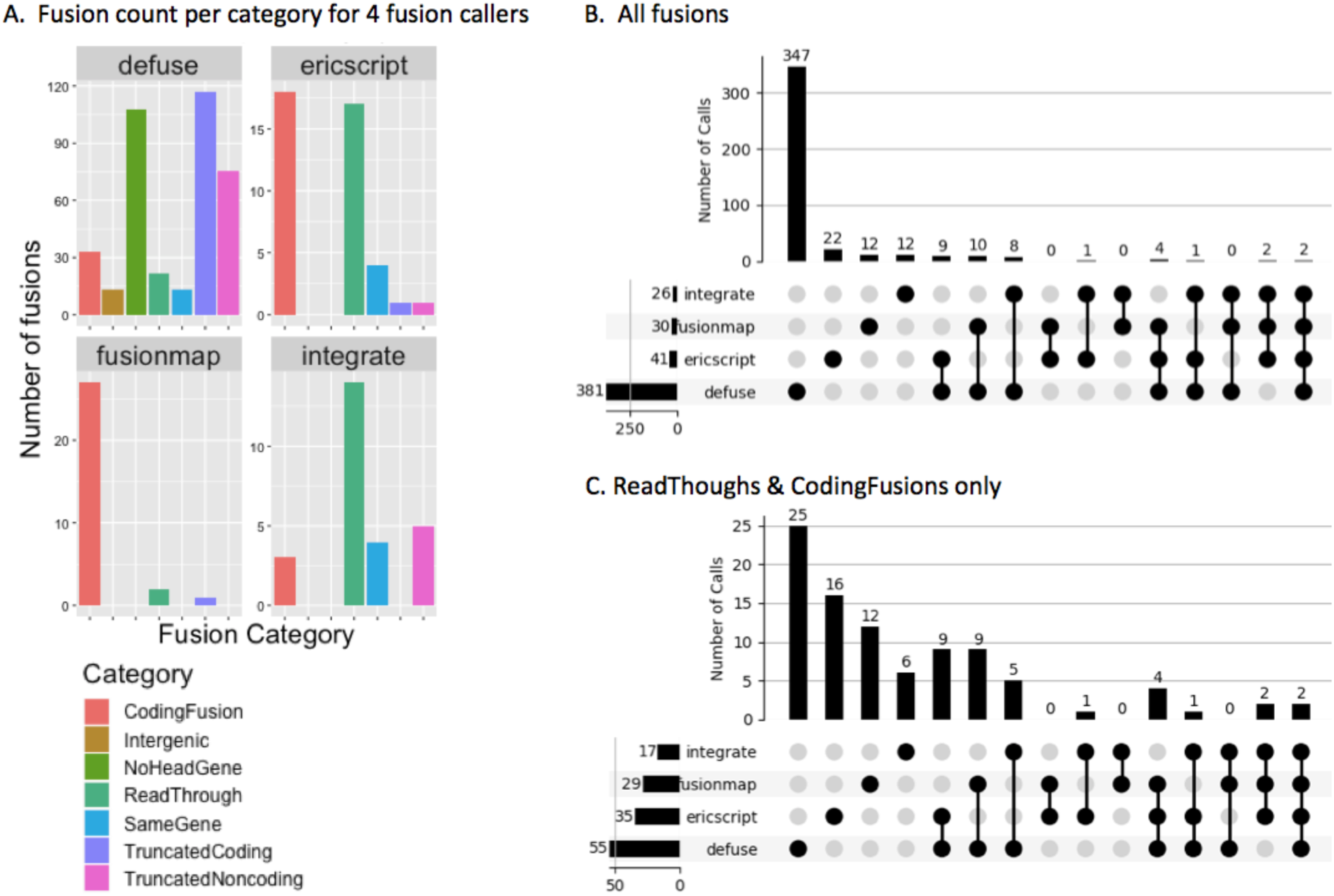
Breakdown of calls made by individual fusion callers. (**A**) Bar graphs showing the number of fusions from each category called by the 4 individual fusion callers used in our pipeline. Category legend is shown below. Not all callers are able to equally identify fusions from all 7 categories. Most of the TruncatedCoding and TruncatedNoncoding fusions were reported by deFuse. FusionMap mainly reports CodingFusions. More than half of the fusions reported by INTEGRATE belong to the ReadThrough category. EricScript primarily reports CodingFusions and ReadThroughs. (**B**) Set intersections for fusions from all 7 categories for the 4 callers used. (**C**) Set intersections for ReadThrough and CodingFusions only for the 4 callers used. All analyses were performed for fusions identified using BWAfilter30, Valfilter3 and Covfilter5 in the total 51 DIPG patient samples.

**Figure 8:**
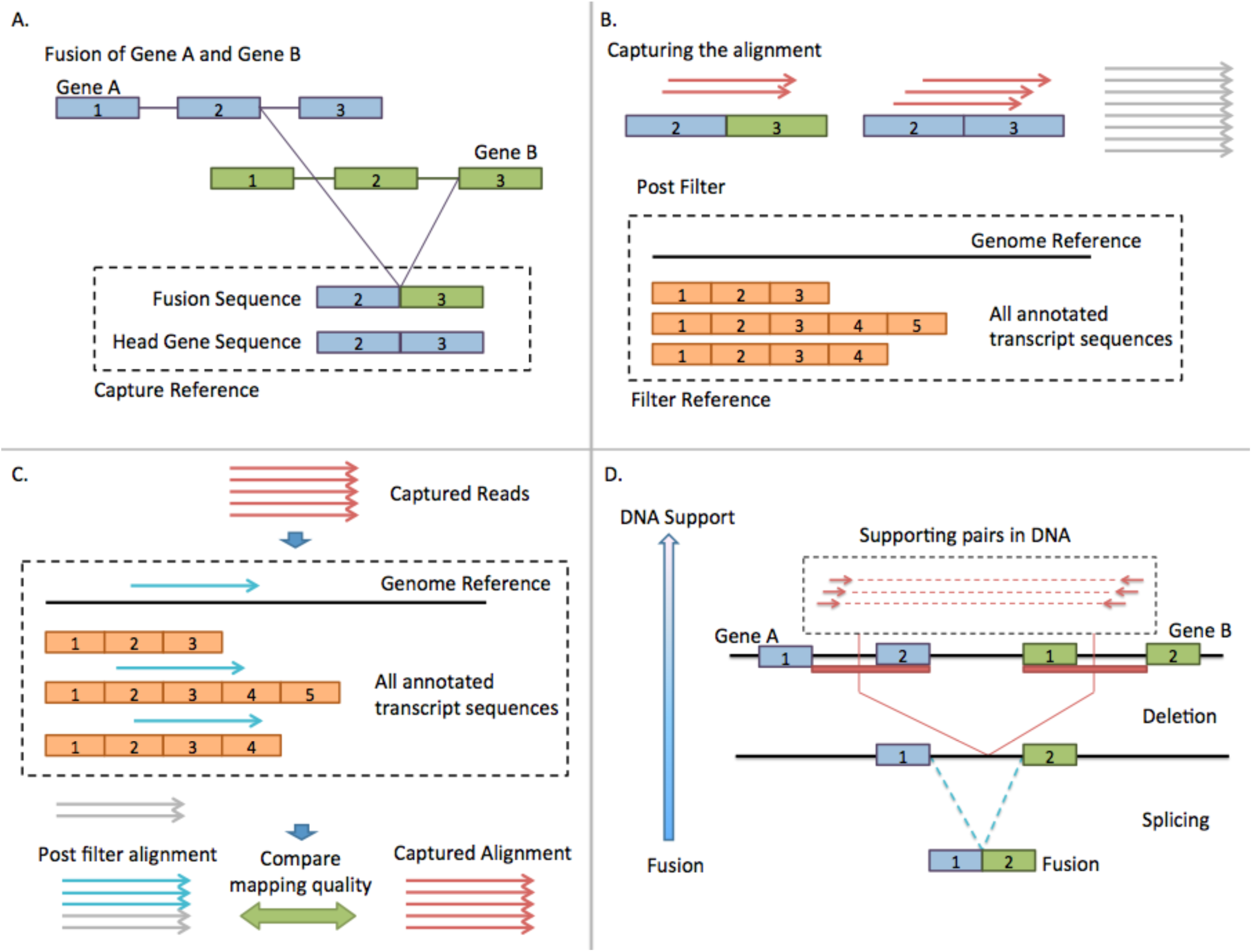
Diagram of the fusion validation pipeline (**A-C**) and DNA support step of the fusion detection pipeline (**D**). (**A**) Creation of 200bp capture mini-contigs for both fusion breakpoint and head gene exon-exon junction sequence. **(B)** Alignment of RNAseq reads to both contigs to allow for the capture of supporting reads (top). Filter reference (bottom). (**C**) Removal of reads which align to filter reference. (**D**) Capture of discordant reads using WGS data, which is indicative of structural variations in DNA.

Seven fusions were unique to DIPG-T samples (Figure 4; Supplementary Table S6), including 2 CodingFusions (*SSBP1-MGAM, NEDD1-CFAP54*), 4 NoHeadGene fusions (*N/A-RP11-152L7.1, N/A-RP11-627D16.1, N/A-CHDC2, N/A-ZNF334*) and a TruncatedNoncoding fusion (*RP11-419K12.1-N/A*). Two NoHeadGene fusions and the TruncatedNoncoding fusion involve long non-coding RNA (lncRNA). Little is known about how lncRNA fusions impact oncogenesis, however they appear to regulate wild-type gene expression in prostate (Qin et al. 2017) and liver (Zhu et al. 2019) cancers. Patients with advanced liver cancer are reported to have more lncRNA fusions compared to those with less severe diagnoses. We also observed that our chosen cohort of 65 fusions generally had more abundant expression in tumours compared to normal sample types across all sample sets. Some fusions were expressed almost ubiquitously across all tumours and normal tissues, but were absent from DIPG-N. These fusions were less abundant in GTEx brain cortex samples, and their absence in DIPG-N may be due to a lack of tissue-specific expression in the frontal lobe. We cannot, however, discount the possibility that this is an artifact of the differences in sequencing used between the various sample sets.

### Intrachromosomal distances between fusion genes in tumours and normal tissues

Our results indicate that fusions in DIPG-T and DIPG-N largely differ in the characteristics of their constituent genes. To determine if these differences extend to gene loci, we compared gene distances for fusions composed of head and tail genes on the same chromosome. We calculated mean and median distances for 2 sets of fusions that passed BWAfilter30, Valfilter3, and Covfilter5 in our 51 unpaired DIPG samples: 1) the 56 intrachromosomal fusions present in both DIPG-T and –N samples, and 2) the 100 DIPG-T-specific intrachromosomal fusions. The mean distance between genes in fusions common to DIPG-T and -N was 16kb, while the median distance was 0kb. The mean and median distances between genes for DIPG-T-specific fusions were substantially larger at 5047.5kb and 7.7kb, respectively, suggesting that fusions unique to tumours tend to occur between genes that are further apart.

### Gene fusions resulting from interchromosomal events

Increased distances between intrachromosomal fusion gene pairs may indicate that broad structural changes are occurring in DIPG to give rise to these observations. We further investigated this possibility by comparing the frequency of fusions between RNA transcripts produced from loci on different chromosomes in normal and tumour tissue samples. For this, we focused on interchromosomal fusions that passed BWAfilter30, Valfilter3, and Covfilter5. The fraction of unique interchromosomal events among DIPG-T-specific fusions was higher than among those common to DIPG-T and DIPG-N samples (Table 2). For the 15 paired DIPG-T/DIPG-N samples, we observed that 5.1% (6/118) of DIPG-T-specific fusions and 4.4% (5/114) of fusions found in DIPG-N were interchromosomal. For the total 51 samples, interchromosomal events occurred in 12.8% (33/256) of DIPG-T-specific fusions and 6.9% (12/174) of fusions in DIPG-N. We also analyzed the dataset published by Wu et al (2014), and found that 15.5% (167/1076) of fusions they reported were interchromosomal. In contrast, 6.9% (49/711) of fusions in 157 brain tissue GTEx samples were interchromosomal.

**Table 2.** Number of inter-chromosomal fusion events in tumour samples, normal samples and both tumour and normal samples across various data sets (across all fusion categories).

|  | Tumour-specific |  | Normal-specific/Tumour&Normal |  |
| --- | --- | --- | --- | --- |
|  | total | Inter-chr (%) | total | Inter-chr (%) |
| **DIPG paired** | 118 | 6 (5.08) | 114 | 5 (4.39) |
| **DIPG all** | 256 | 33 (12.90) | 174 | 12 (6.89) |
| **Wu et al 2014** | 1076 | 167 (15.52) | -- | -- |
| **GTEx** | -- | -- | 711 | 49 (6.89%) |

### Evaluating underlying DNA Support for fusions

Of our 51 DIPG samples, 14 DIPG-T and 13 DIPG-N had both WGS and RNAseq data available, allowing us to compare the presence of fusions between their genomes and transcriptomes. We examined the WGS data for evidence of fusions previously identified in RNAseq data using BWAfilter30, Covfilter5 and Valfilter3 (Table 3). The WGS data for each sample was sequenced to 1495 million average number of reads per sample (max=2206 million; min=1021 million). We identified DNA support for 5 CodingFusions (*ERC1-FAM222B, LDLRAD3-NCAM1, TNRC6B-CELF2, NMU-PDGFRA*, and *WDFY3-FAM471-STBD1*) and 1 TruncatedCoding (*DDX1-MYCNUN*) (Table 3). All 6 of these fusions were specific to DIPG-T samples. While DNA restructuring for ReadThrough fusions is not expected, as they are typically caused by transcriptional read-through and *cis*-splicing of the resultant chimeric transcripts (Qin et al. 2015; Tang et al. 2017), we also observed no evidence of genomic restructuring for TruncatedNoncoding, NoHeadGene or SameGene fusions, which are all typically attributed to chromosomal translocations.

**Table 3.** Fusion numbers of samples with WGS data for 14 DIPG-T and 13 DIPG-N samples using BWAfilter30, Valfilter3 and Covfilter5. Number of fusions with DNA support is shown in brackets.

| Category | DIPG-T | DIPG-N | Both |
| --- | --- | --- | --- |
| ReadThrough | 18 | 1 | 4 |
| CodingFusion | 20 (5) | 0 | 2 |
| TruncatedCoding | 47 (1) | 6 | 18 |
| TruncatedNoncoding | 20 | 9 | 20 |
| NoHeadGene | 31 | 13 | 20 |
| SameGene | 4 | 0 | 9 |
| **Total** | **140 (6)** | **29 (0)** | **73 (0)** |

We also performed Sanger sequencing and a series of PCRs to experimentally validate several fusions either with or without WGS/WES support. First, we selected 4 of the above 6 fusions with WGS/WES DNA support as well as 4 fusions with DNA support that did not pass BWAfilter30, Covfilter5 or Valfilter3 (Supplementary Table S7). We performed genomic PCRs for 7 out of 8 fusions and successfully identified them in the DNA. Using RT-PCR, we were also able to identify all 8 of these fusions in the RNA. We then selected 8 RNA-specific fusions, 3 of which passed all filters and 5 which failed at least one (Supplementary Table S8). RNA transcripts for all 8 of these fusions were successfully identified using RT-PCR. Sanger sequencing was used to confirm the head and tail genes of each PCR product.

### Increased expression of spliceosomal genes correlates with fusion occurrence

Our initial WGS findings suggest that a majority of fusions in DIPG occur only in RNA. RNA-specific fusions have been previously reported and can result from various splicing aberrancies (Li et al. 2008; Qin et al. 2015). We thus examined the expression of genes related to splicing activity and spliceosomal function in DIPG-T patients. From the RNAseq data we estimated gene expression levels in each DIPG-T sample and used this to cluster the expression of genes in the Kyoto Encyclopedia of Genes and Genomes (KEGG) spliceosome category (Kanehisa and Goto 2000). Samples fell into 1 of 2 broad groups with either increased or decreased spliceosomal gene expression (Figure 5). Samples with lower spliceosomal expression also had fewer total fusions than samples with higher spliceosomal expression (mean fusion number 40.5 and 57.3; p=0.007, Wilcoxon test), suggesting a link between expression of spliceosomal genes and occurrence of fusions in the DIPG tumour samples.

#### Comparison of expression levels and fusion counts

Distributions of raw fusion counts are different between DIPG-T and DIPG-N samples (Figure 3C, Figure 5). DIPG-T samples have a much broader range of fusion counts than DIPG-N samples, which are generally more consistent across the sample cohort. Fusion counts also correlated positively with spliceosomal gene expression within DIPG-T samples (Figure 5). DIPG28T, for example, has the highest fusion count among paired samples and one of the highest expression level profiles for spliceosomal genes. It likewise shows a marked increase in fusion number compared to its paired normal DIPG28N (Figure 3). DIPG25T, on the other hand, has an extremely low fusion count (Figure 3) and exhibits the greatest reduction in spliceosomal gene expression (Figure 5). It also has a substantially lower fusion count than its paired normal sample, DIPG25N (Figure 3). DIPG71T has a relatively low number of fusions in comparison to DIPG71N, and generally under-expresses spliceosomal genes. Similar trends can be seen with DIPG14BT/N.

#### NEDD1-CFAP54 is a fusion with characteristics of aberrant splicing without support in genomic DNA

Our results show that fusion formation in DIPG is complex and can be independent of chromosomal restructuring. This is exemplified by the DIPG-specific fusion between the neural precursor cell expressed, developmentally downregulated 1 (*NEDD1*) gene and cilia and flagella-associated protein 54 (*CFAP54*) gene, which is also known as *C12orf55*. Our pipeline identified supporting reads for *NEDD1-CFAP54* in the transcriptomes of 4/34 DIPG-T samples with no evidence of corresponding DNA rearrangements. We also confirmed the presence of this fusion in the RNA of 3/4 samples using RT-PCR and Sanger sequencing (Supplementary Figure S3). Interestingly, *CFAP54* is immediately upstream of *NEDD1* on chromosome 12 (12q23.1), yet the fusion contains *NEDD1* as the head gene and *CFAP54* as the tail gene. The same breakpoint in *NEDD1* is consistent across all *NEDD1-CFAP54*-positive samples, whereas two breakpoints are present in *CFAP54* – one that is ubiquitous and one in a second variant of *NEDD1-CFAP54* in DIPG89T. We also observed several instances of a DIPG-T-specific *CFAP54-NEDD1* ReadThrough with varying breakpoint combinations in both genes, although the frequency of this fusion orientation was lower than that of *NEDD1-CFAP54*. Samples DIPG52T, DIPG62T and DIPG89T contained both the *NEDD1-CFAP54* and *CFAP54-NEDD1* fusions, potentially indicating the formation of a circular RNA. Of note, these samples all show increased spliceosomal gene expression and fusion counts, with DIPG89T having the highest total fusion number of all samples. While the exact processes remain to be determined, these results suggest that dysregulated spliceosomal expression and RNA-specific fusion formation are interrelated.

## Discussion

Gene fusions are causal to several cancers, and understanding their effects and formation is critical for understanding cancer pathology. Most fusion-calling methods, however, are hindered by the ways in which they define what constitutes a true gene fusion. We have thus developed a pipeline that consolidates information from 4 fusion callers with varied definitions, allowing for a greater capture of fusions in a given sample set. This pipeline also examines WGS data to determine if DNA support is present for fusions found in RNAseq. The pipeline is designed to be easily updated with additional fusion calling methods, making it versatile and applicable to research beyond the scope of this paper.

We used our pipeline to compare tumours and normal brain tissue samples from DIPG patients, revealing that although these tissues have a comparable total number of fusions per sample, DIPG-T samples have a bias for non-recurrent fusions and fusions between coding genes. An increase in sample-specific fusions is not surprising; cancer genomes harbour more instability, and thus contain more translocations and fusion events, many of which are thought to be non-recurrent (Bunting and Nussenzweig 2013; Kim and Jinks-Robertson 2012; Yoshihara et al. 2015). Furthermore, both intra- and inter-tumour heterogeneity is well-documented in various cancers, including gliomas (Abou-El-Ardat et al. 2017; Alizadeh et al. 2000; Aum et al. 2014). We also show that ReadThroughs and CodingFusions were almost exclusively found in DIPG-T (Figure 3A, B). ReadThroughs typically result from *cis*-splicing, in which two neighbouring genes are transcribed into a single pre-mRNA (Qin et al. 2015; Tang et al. 2017) and, thus, typically do not have DNA support. CodingFusions, on the other hand, consist of non-adjacent genes and, unlike ReadThroughs, they cannot be accounted for by transcriptional read-through and *cis-*splicing. Instead, they are typically attributed to chromosomal rearrangements or translocations. In a recent study, nearly half of the paediatric DIPG patients examined had fusion-forming structural variants in their DNA (Wu et al. 2014), suggesting that chromosomal translocation is relatively common in DIPG.

Surprisingly, we found DNA support for only 6 of 242 fusions for which genomic data was available (Table 3). Fusions specific to RNA are increasingly reported in cancers and normal cell lines (Babiceanu et al. 2016; Grosso et al. 2015; Jia et al. 2016; Li et al. 2008; Qin et al. 2015; Yun et al. 2014). Similarly, in our study we have not been able to identify DNA support for most fusion events, suggesting they appear exclusively in the transcriptome. This lack of DNA support indicates that these fusions are likely driven by dysregulation of RNA transcription, splicing or processing. Multiple breakpoint combinations and gene orientations in individual samples, such as those in fusions involving the *NEDD1* and *CFAP54* genes, likewise point to impaired RNA mechanisms.

In keeping with a dysregulation of RNA mechanisms, DIPG-T samples generally over- or under-expressed genes in the KEGG spliceosomal category, and showed a positive correlation between spliceosomal gene expression and fusion number (Figure 5). Co-occurrence of gene fusions and splicing gene mutations has been previously reported in mesothelioma (Bueno et al. 2016), and correlations between splicing factor dysregulation and fusion expression have been established in prostate cancer (Qin et al. 2015). In DIPG, increased spliceosomal gene expression may result in over-activation of splicing mechanisms to give rise to an accumulation of RNA fusions. ReadThroughs form via *cis*-splicing, while *trans*-splicing of two separately transcribed pre-mRNA molecules can drive RNA-specific formation of fusions typically thought to be caused by genomic restructuring, such as CodingFusions (Gingeras 2009; Li et al. 2008). *Cis*-splicing, *trans*-splicing and splicing to produce truncated- and NoHeadGene-like fusions have all been previously reported in various cancers (Bartonicek et al. 2017; Li et al. 2008; Jia et al. 2016; Qin et al. 2015). While the relationship between spliceosomal gene dysregulation and the occurrence of fusions in DIPG remains unclear, our results point to mechanisms of fusion formation beyond chromosomal rearrangement that should be taken into account when investigating potential routes to oncogenic transformation in cancer in general.

## Materials and methods

### Sample cohorts

#### DIPG patient tissue samples

Our study cohort comprised of 51 samples from 36 DIPG patients. We obtained 34 samples from DIPG tumour biopsies (DIPG-T) and 17 normal brain tissue samples from autopsy of the frontal lobe (DIPG-N). Of these, 30 were DIPG-N/DIPG-T paired samples obtained from 15 patients. The remaining 19 DIPG-T and 2 DIPG-N samples were singletons. We excluded corresponding DIPG-T data for these 2 DIPG-N samples due to low RNAseq quality. All samples have 100bp paired-end RNAseq data. 27 samples (14 DIPG-T, 13 DIPG-N) have 100bp paired-end WGS data. Total RNA was extracted from tissues using an RNeasy mini kit (Qiagen).

#### Additional tumour and normal tissue samples

We obtained RNAseq datasets for tumour and corresponding normal tissue samples from the TCGA, totalling 283 samples. We selected samples from breast cancer (BRCA; 20 tumour, 20 normal), colon adenocarcinoma (COAD; 20 tumour, 20 normal), glioblastoma multiforme (GBM; 20 tumour, 5 normal), kidney renal clear cell carcinoma (KIRC; 20 tumour, 20 normal), brain lower grade glioma (LGG; 20 tumour), liver hepatocellular carcinoma (LIHC; 19 tumour, 20 normal), prostate adenocarcinoma (PRAD; 20 tumour, 20 normal), and thyroid carcinoma (THCA; 19 tumour, 20 normal). See Supplementary Table S9 for complete list of TCGA samples. From GTEx, we obtained RNAseq data of a total 1277 samples from 53 normal tissue types (Supplementary Table S10). Finally, we utilize a validation dataset composed of RNAseq data from 73 glioma (DIPG and HGG) samples (Wu et al. 2014).

### Development of a gene fusion detection pipeline

#### Overview of the gene fusion detection pipeline

To systematically identify a broader range of fusions from RNAseq data, we developed a detection pipeline that incorporates multiple fusion calling approaches. We incorporated deFuse (McPherson et al. 2011), FusionMap (Ge et al. 2011), Ericscript (Benelli et al. 2012) and INTEGRATE (Zhang et al. 2016) into our pipeline, which was developed under the GenPipes framework (Bourgey et al. 2019), as shown in Figure 6. Each fusion caller runs on the input data to identify fusions independently. The gene names, breakpoints and strand information are taken from each caller’s output file and converted into a Common Fusion Format (CFF) --which unifies the file formats of each caller into a single, consistent output -- and re-annotated using custom annotation scripts. The fusion calls are then assigned to 1 of 7 categories (Figure 2 and see below). Fusions are then checked for evidence of DNA support using available genomic data. Next, fusions that map to low mappability regions and fusions with fewer than 3 supporting split reads from our validation pipeline are removed (BWAfilter30 and Valfilter3, respectively; see below). Fusion calls are then merged/clustered based on similarity of head/tail genes and proximity of their breakpoints. Finally, merged entries are kept only if they have both 5 split and 5 spanning reads (Covfilter5). Since our pipeline uses a Common Fusion Format, it is straight-forward to incorporate additional callers as they become available.

#### Consistent categorization of fusions

Gene fusions are classified into 7 broad categories according to the features of the constituent genes and the breakpoint loci (Figure 2). For a given fusion, we refer to the upstream gene the “head gene” and the downstream gene the “tail gene.” When both genes are coding and the head gene is immediately upstream of the tail gene, the fusion is categorized as a “ReadThrough.” If both genes are coding and are either on different chromosomes or on the same chromosome but not in a ReadThrough orientation, the fusion is categorized as a “CodingFusion.” In cases where the head gene is coding and the tail gene is non-coding or absent, such as if the downstream breakpoint falls into an intergenic region, the fusion is assigned to the “TruncatedCoding” category. Similarly, when the head gene is non-coding, the fusion is assigned to the “TruncatedNonCoding” category. It should be noted that TruncatedCoding and TruncatedNoncoding fusions likely undergo nonsense-mediated decay before translation can occur (Mendell et al. 2004). Fusions with no meaningful head genes, such as those in which the upstream breakpoint can only be mapped to intergenic region, are assigned to the “NoHeadGene” category (Figure 2). In some cases, the two breakpoints of a fusion can be mapped to two different genes as well as to a longer gene overlapping those two genes. Such fusions are assigned to the “SameGene” category.

#### Establishing fusion filters for the detection pipeline

Though the four fusion callers incorporated into our detection pipeline all have their own filters, we added additional filters in downstream analyses to increase confidence in the result we obtained. We use 3 filters in our pipeline: BWAfilter30, Valfilter3 and Covfilter5.

The BWAfilter30 is used to check the uniqueness, or “mappability” of boundary sequences surrounding fusion breakpoints. Fusions picked up due to non-unique boundary sequences are more likely to be false positives, and are therefore removed. We extract 30 nucleotides upstream and 30 nucleotides downstream of a given breakpoint, and then map these 60 nucleotide contigs to the human genome using the Burrows-Wheeler Aligner (BWA). Fusions with boundary sequences that are found to be less than 100% unique are then removed by this filter.

Next, we use our validation pipeline (Valfilter3) to directly check for supporting split reads in a sample’s RNA-seq data (see Methods below and Figure 8 for details). Only calls for which our validation pipeline captures 3 or more split reads are kept. Additionally, our validation pipeline can identify fusions that were not called by the detection pipeline. This may occur, for example, in instances where a fusion is present in a given sample, but is expressed at a level that is too low to pass the detection pipeline filtering steps. We use this feature to obtain a more accurate number of the samples a given fusion is present in. Following this step, calls from multiple callers that correspond to the same fusion event are merged into a single entry.

After merging calls together, we set a coverage filter of at least 5 split reads and 5 pair/spanning reads (Covfilter5). Higher read coverage for a fusion indicates it is more likely to be true. FusionMap does not provide spanning read counts, so we only applied Covfilter5 to split reads for this caller.

### Validation dataset creation

To evaluate the accuracy of the fusion detection pipeline, we used a previously published dataset of 75 glioma samples (31 DIPGs and 44 HGGs) (Wu et al. 2014). Two of the samples (SJHGG139_D and SJHGG141_D) were excluded due to damaged data files, leaving 73 samples in our final validation set. The dataset contained 144 chromosomal structural variants (SVs) that were reported to result in the formation of fusions. From these we removed the SVs that were not validated, whose breakpoints could not be mapped to genes, or who were present in samples SJHGG139_D and SJHGG141_D. After filtering, reannotation and merging by our pipeline, a validation set of 104 fusions remained.

### Assessing fusion caller preference for fusion categories

We applied our fusion detection pipeline to all DIPG-T and DIPG-N samples. First, we used each of our 4 chosen fusion callers to identify fusions in our sample set, and then categorized the fusions (Figure 7A). In total, 430 fusions were reported by the combined results of the 4 callers for all categories, and 92 for ReadThrough/CodingFusion only. We then used the BWAfilter30 and Covfilter5 fusion filters and compared the overlap in the number of fusions reported by each caller (Figure 7B). With this, we show that the 4 callers do not predict fusion categories equally.

### Developing a fusion validation pipeline

#### Overview of the fusion validation pipeline

The validation pipeline checks whether a given fusion exists in an RNAseq dataset by looking for supporting split reads for that fusion (Figure 8A-C). It can function as part of the detection pipeline (Valfilter) or independently. To query for a fusion, we extract 100bp from either side of the breakpoint to build a “capture reference” (Figure 8A), and then align all reads to this reference in order to capture potential supporting reads (Figure 8B). Reads that successfully align to the “capture reference” are termed “captured reads”. To ensure captured reads are not from other regions of the genome or from junctions of exons, we build a “filter reference”, comprising of whole genome sequences and all transcriptome sequences based on our combined annotation (ensembl genes plus known genes). We align captured reads to this filter reference and remove captured reads which align better to the filter reference (Figure 8C). Reads remaining after filtering are considered “supporting reads” for fusions. In the capture alignment (Figure 8B), the read set is large and reference set small (only fusion sequences). In the filter alignment (Figure 8C), the reference set is large (consisting of the whole human genome plus the transcriptome), and read set small (consisting of only the filtered read set). This enables the two steps to have better running time than fusion caller tools, which require aligning of the full read set to the whole genome or transcriptome. Thus, our fusion validation pipeline is much faster than our fusion detection pipeline. The fusion validation pipeline is also more sensitive than the fusion detection pipeline, as it can validate fusions in samples where they were not found initially. This fusion validation pipeline is able to query the transcriptome for provided fusions, but is unable to call fusions from scratch like fusion callers do.

### Identification of DNA support for known fusions

Gene fusions caused by structural variants in the genome, such as deletions, inversions or translocations, can be expected to have supporting read pairs in WGS data. Thus, for samples with available WGS data, we search for read pairs supporting the fusions identified in those samples. We searched for pairs in which 1 read mapped to the region from the upstream breakpoint to the end of head gene and the other read mapped in the region from the start of tail gene to the downstream breakpoint (Figure 8D). We clustered these pairs with a minimal requirement of 3 pairs, and considered these clusters as DNA support for fusions.

### Experimental validation of the fusion pipeline by PCR

In order to validate our pipeline and the fusions identified, we selected 16 fusions from the 7 fusion categories in our DIPG samples (Supplementary Table S6). Seven of these fusions passed filtering using BWAfilter30, Valfilter3 and Covfilter5. The remaining 9 fusions failed either one or two filters yet had some interesting features, such as being recurrent, having both breakpoints on exon boundaries, or having DNA support. Of the 16 fusions, 8/16 were CodingFusion category fusions. Using PCR, we validated the presence of all 16 fusions in RNA and 7/16 in DNA.

Total RNA from normal and tumorous brain tissues was isolated by using a Qiagen RNeasy Mini Kit and quantified on a NanoDrop spectrophotometer (Thermo Fisher Scientific). 1 μg of each RNA sample was converted into single-strand cDNA with random hexamers using an iScript™ Select cDNA Synthesis Kit (Bio-Rad). To validate fusion transcripts, regular PCR assays were carried out on a SimpliAmp™ Thermal Cycler (Thermo Fisher Scientific) with a final volume of 50 μl reaction mixture containing cDNA template, 25 μl HotStarTaq Master Mix (Qiagen) and 10 pmol of each primer. The PCR cycling conditions were an initial “Hot Start” activation step at 95°C for 15 min, followed by 40 cycles of 30 s at 94°C, 30 s at 60°C and 1 min at 72°C, and a final extension at 72°C for 10 min. The sequences of all primer pairs are listed in Supplementary Table S7 and S8. The housekeeping β-actin gene was amplified for cDNA quality control and a non-template negative control was included in each PCR run. PCR products were separated on 1% agarose gel, purified with a QIAquick Gel Extraction kit (Qiagen) and sequenced in both directions with the original set of primers on a 3730XL DNA Analyzer (Applied Biosystems) at the Sanger Sequencing Facility of the Centre for Applied Genomics, the Hospital for Sick Children. Sequences were then compared with the fusion transcripts identified by RNA-Seq by using the BLAST program from NCBI (http://blast.ncbi.nlm.nih.gov/Blast.cgi).

### Gene expression estimates

Raw sequences were mapped to the GRCh37 version of the human genome and transcriptome using STAR (version 2.5.0c) (Dobin et al. 2013). The resulting bam files were merged based on sample identities, sorted by coordinates and marked for duplicates using Picard Tools (version 1.123; https://broadinstitute.github.io/picard/). The final bam files were used to estimate gene expression (FPKM) using Cufflinks (version 2.2.1) (Trapnell et al. 2012).

### Clustering of the data

We used the R package “pheatmap” for clustering the data based on average linkage and euclidean distance for both genes and samples using the FPKM values estimated using Cufflinks.

### Construction of the “filter reference” for the validation pipeline

The filter reference is a .fasta file consisting of the whole hg19 genome reference and transcript junction sequences. To build these transcript junction sequences, we use the function “build_junction_seq_for_gene_bed(ref, ensbed)” in our custom annotation pipeline “pygeneann.py”. The “ref” is the hg19 genome reference, and the “ensbed” is a .bed format file genomic feature file containing ENSEMBL genes and known genes, which was generated using the Table browser feature of the UCSC genome browser. The filter reference, ensbed and ref files are provided as part of our pipeline.

## Data access

All RNAseq data generated from this study will be submitted to the Gene Expression Omnibus (GEO; https://www.ncbi.nlm.nih.gov/geo/) upon acceptance of the manuscript.

## Acknowledgements

Bioinformatic analyses were supported by the Canadian Center for Computational Genomics (C3G), part of the Genome Technology Platform (GTP), funded by Genome Canada through Genome Quebec and Ontario Genomics.

